# Interrogation of retinal lipofuscin by fluorescence lifetime imaging microscopy

**DOI:** 10.1101/2025.09.12.675809

**Authors:** V. Andreev, M. Yakovleva, A. Kostyukov, V. Sokolova, V. Shcheslavskiy, G. Goltsman, T. Feldman, V. Kuzmin, M. Ostrovsky, P. Morozov

## Abstract

Age-related macular degeneration is a disease that affects the middle part of the vision and involves pathological alterations in the retinal pigment epithelium. Accurate and timely evaluation of the retinal pigment epithelium is a cornerstone of effective treatment planning. In this study, we present the development of a preclinical method for early diagnostics of age-related macular degeneration using time and spectral characteristics of fluorescence of lipofuscin granules from retinal pigment epithelium. Using the unique system based on asuperconducting single-photon detector and time-correlated single-photon counting electronics integrated in the confocal laser scanning microscope we determined the parameters of fluorescence (distribution long and short fluorescence lifetime components and their contribution to the total fluorescence signal as well as fluorescence spectral shift) that have a diagnostic value for differentiation of the normal and pathological states in the degenerative diseases of the retina and retinal pigment epithelium.

## Introduction

Age-related macular degeneration (AMD) is one of the leading causes of vision loss in the elderly [1-3]. The progression of AMD involves pathological alterations in the retinal pigment epithelium (RPE), ultimately resulting in the degeneration and death of photoreceptor cells [4]. Currently, AMD is considered incurable, making early detection critical for implementing timely protective and preventive measures. Such interventions can slow disease progression, helping patients maintain a good quality of life for years.

In light of this, researchers are actively developing novel methods and refining existing techniques for the preclinical diagnosis of retinal and RPE disorders. Among these, fundus autofluorescence (FAF) imaging has emerged as a particularly promising non-invasive approach [5,6]. FAF enables detailed visualization of pathological changes in the fundus, allowing for the study of early AMD manifestations and disease dynamics.

FAF imaging enables clinicians to evaluate RPE cell health in AMD patients, guide treatment decisions, and predict disease progression. As one of the most sensitive modern techniques for detecting early retinal/RPE changes [7], FAF is crucial for identifying AMD in its initial stages. Moreover, it offers an objective, high-resolution assessment of treatment efficacy.

However, a key limitation of FAF imaging is its inability to provide quantitative measurements of pathological changes. While FAF pattern analysis yields detailed qualitative data—allowing clinicians to detect abnormal zones and distinguish between different pathological forms—it relies on subjective interpretation. Disease progression is assessed by comparing a patient’s FAF pattern with reference images from healthy individuals, which introduces variability. Additionally, FAF cannot detect pathology at the preclinical stage, when symptoms are absent and the autofluorescence pattern appears normal.

Given these constraints, current research highlights the need to further refine FAF imaging to enhance its diagnostic precision, thereby improving early AMD detection and prognostic accuracy.

FAF imaging primarily detects lipofuscin granules (LGs), commonly known as the “aging pigment.” These granules accumulate in retinal pigment epithelium (RPE) cells as a result of incomplete lysosomal degradation of phagocytosed photoreceptor outer segments. This process is significantly accelerated in retinal and RPE degenerative diseases.

The fluorescent properties of LGs originate from *all-trans*-retinal conjugates, primarily bis-retinoids (Bis-Rets) such as A2E, along with their photooxidation and photodegradation products (Oxy-Bis-Rets) [8, 9]. These compounds define the spectral and kinetic characteristics of autofluorescence and may serve as biomarkers for pathological changes in AMD.

LGs are photosensitive and, upon absorbing visible light, generate reactive oxygen species (ROS) [10]. Bis-Rets are the primary source of ROS in LGs, but they are also susceptible to oxidation upon ROS interaction, forming Oxy-Bis-Rets. The fluorescence spectrum of these oxidized products shifts toward shorter (blue) wavelengths [11]. Oxy-Bis-Rets exhibit high toxicity and can accumulate in RPE cells, damaging cellular structures even in the absence of light [12–14]. Studies have shown elevated Oxy-Bis-Rets levels in AMD compared to normal conditions [15, 16], suggesting that their increased presence—and the resulting changes in spectral-kinetic properties could serve as an early diagnostic marker [16, 17].

Both qualitative and quantitative assessments of LG fluorescence (spectral properties and decay kinetics) could provide crucial prognostic insights into retinal degenerative diseases.

A particularly promising advancement involves measuring fluorescence lifetime at specific wavelengths [18]. *In vivo* studies have revealed differences in fluorescence lifetime between healthy retinas and those with AMD [19]. These measurements were obtained using a fluorescence lifetime imaging ophthalmoscope (FLIO), adapted from a confocal laser scanning ophthalmoscope (Heidelberg, Germany), which delivers highly reproducible FAF lifetime data in the macular region. Furthermore, fluorescence parameters in healthy tissues differ significantly from those in patients with diabetes mellitus [19], glaucoma [20], and Alzheimer’s disease [21]. Despite progress, precise diagnostic criteria distinguishing normal from pathological states via fluorescence lifetime imaging microscopy (FLIM) remain undefined, limiting its clinical adoption in ophthalmology.

Previous studies analyzing LG fluorescence decay using time-correlated single-photon counting (TCSPC) approach demonstrated that Oxy-Bis-Rets exhibit longer fluorescence lifetimes than non-oxidized Bis-Rets[15, 22, 23]. This finding explains the observed increase in average fluorescence lifetime in AMD, suggesting that elevated Oxy-Bis-Rets levels in RPE LGs contribute to fluorescence decay in pathology [16, 20, 26]. However, these studies relied on fluorescence lifetime data from point measurements in RPE cell suspensions or chloroform-extracted LGs, which is far away from in *vivo* conditions [24, 25].

Fluorescence lifetime distribution across investigated samples provides significantly more diagnostic information than single-point measurements. FLIM microscopy enables detailed spatial analysis of fluorescence lifetimes in LGs before and after oxidation. This technique can reveal intricate cellular and tissue structural and functional details, allowing for in-depth mapping of fluorescence lifetimes.

In this work, we report an investigation of fluorescence decay properties of LGs for the evaluation ofthe contributions of Bis-Rets and Oxy-Bis-Rets to the overall fluorescence decay signal. Using a unique system based on the confocal laser scanning microscope equipped with a superconducting nanowire single-photon detector (SSPD) and time-correlated single photon counting electronics we determined the parameters that have a diagnostic value for differentiation of the normal and pathological states in the degenerative diseases of the retina and retinal pigment epithelium.

## Materials and methods

### Isolation of LGs

Human cadaver eyes were obtained from the Eye Tissue Bank of the S.N. Fedorov National Medical Research Center for Scientific and Technical Complex “Microsurgery of the Eye” from donors from the thanatology departments of the Moscow Bureau of Forensic Medicine based of the current agreement between the Moscow Bureau of Forensic Medicine and the S.N. Fedorov National Medical Research Center for Scientific and Technical Complex “Microsurgery of the Eye”, as well as the agreement on scientific cooperation between the IBCPh RAS and the S.N. Fedorov National Medical Research Center for Scientific and Technical Complex “Microsurgery of the Eye” [32]. Human cadaver eyes were obtained within 10 hours after the donor’s death following corneal removal for transplantation. Each cadaver eye was autopsied by an ophthalmologist. After removal of the lens, vitreous body and retina, a detailed description of the fundus was performed. Analysis and screening of the donor material were conducted based on clinical signs. The samples were examined under subdued lighting.

LGs were isolated from the RPE of 100 cadaveric eyes of donors aged 50–75 years without any signs of pathology according to the method described elsewhere and suspended in a solution of 0.1 M K-phosphate buffer, pH=7.3 [10]. The concentration of granules was determined by the standard method in a Goryaev chamber. The initial concentration of granules was 3×10^8^ granules/ml.

### Preparation of LG chloroform extracts

Bis-Rets and Oxy-Bis-Rets were extracted from LGs using a chloroform–methanol mixture (1:1) according to Folch protocol [28]. A two-fold excess of a chloroform:methanol mixture (2:1 v/v) was added to the LG suspension. The mixture was stirred using an electric stirrer for 2 min and incubated for 10 min at 4°C. The mixture was centrifuged for 10 min at 4°C. The lower chloroform phase was collected with a syringe, transferred to a flask and evaporated using a vacuum pump (Vacuubrand MZ 2CNT + AK + M + D, Germany). For further chromatographic analysis, each dried sample was re-suspended in 200 μl of methanol.

### High-performance liquid chromatography (HPLC)

Chromatographic separation of Bis-Rets and Oxy-Bis-Rets in chloroform extracts of LGs was performed on a Knauer chromatograph (Germany) with a Kromasil-100-5-C18 column (4 × 250 mm, sorbent size 5 μm). Separation was performed by linear gradient elution in the system: from 80% acetonitrile + 20% water (+ 0.05% trifluoroacetic acid) to 100% acetonitrile in 20 min; flow rate 1.0 ml/min^25^. The products of chromatographic separation were measured using a Knauer K-2501 photometric detector.

### Fluorescence spectra and lifetime measurements

Fluorescence spectra were recorded using an RF-5301 PC fluorimeter (Shimadzu, Kyoto, Japan) equipped with an R955 photomultiplier detector (Hamamatsu, Shizuoka, Japan). RFPC software version 2.0 (Shimadzu) was used for data compilation. Fluorescence was excited at a wavelength of 488 nm with a sampling interval of 1 nm. Fluorescence spectra were corrected for excitation intensity using the spectral response (quantum efficiency) of the R955 photomultiplier detector. All fluorescence spectra were normalized to a wavelength of 592 nm.

The fluorescence lifetime of the suspension of LGs was measured by a FluoTime-300 fluorimeter (PicoQuant, GmbH, Germany). Fluorescence excitation of the sample was performed using an LDH-PC-485 diode laser (485 nm, pulse duration 107 ps). The fluorescence signal was recorded at a wavelength of 540 nm. The fluorescence decays were fitted using a three-exponential model. Fluorescence lifetimes and the contributions of the individual fluorophores to the detected fluorescence signal were evaluated using the formula taking into account the instrument response function (IRF):

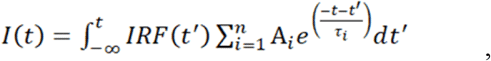

where i is the component number, A is the amplitude, τ is the fluorescence lifetime. The mean fluorescence lifetime was calculated using the formula:

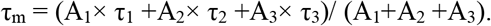

### Synthesis of A2E

Synthesized A2E was used as a standard. A2E was obtained from *all*-*trans-*retinal and ethanolamine in acetic acid and ethanol as described elsewhere [34]. The purity of A2E was controlled by HPLC on a Knauer chromatograph (Germany). A2E was identified using a 7T LTQ FT mass spectrometer (Thermo Electron Corp., Germany) equipped with an electrospray ion source. Mass spectra were processed and analyzed using Qual Browser 1.4.

### Irradiation of samples

Photooxidized LGs (AMD model) were obtained by irradiating LG samples with visible light (400‒700 nm) using a Led15w – 4000K lamp (the surface density of the light energy flux incident on the sample was 100 mW/cm^2^).

### Fluorescence lifetime imaging microscopy with a superconducting single-photon detector

FLIM was performed using Zeiss Axio Observer A1 microscope (Carl Zeiss, Germany) equipped with a confocal DCS-120 laser scanner and photon counting electronics (SPC-150NX) (Becker&Hickl, Germany). The optical system of the scanner was modified to allow coupling of the optical fiber from a superconducting single-photon detector (Fig. 1).

**Fig. 1.**
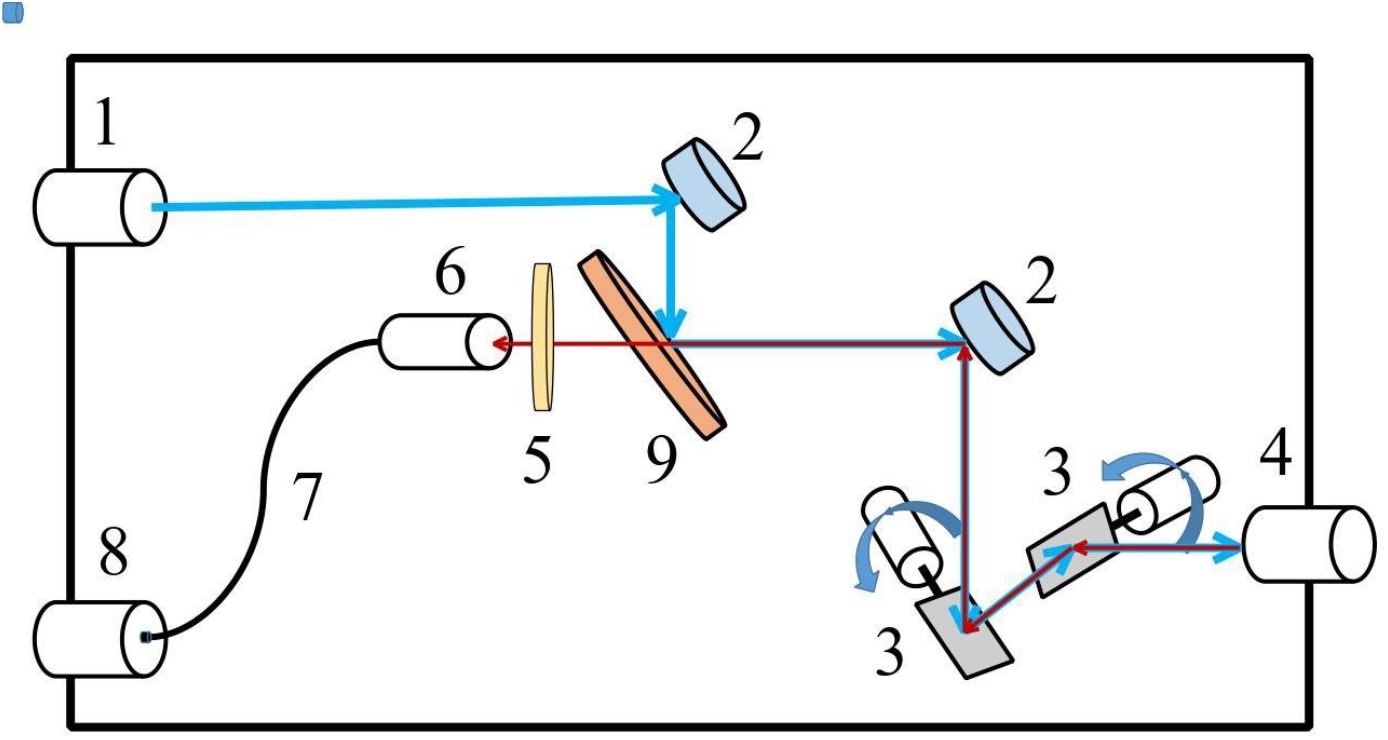
Optical scheme of the setup. 1 – Laser-in port, 2 – Mirror, 3 – galvanometric mirrors, 4 – scanning lens, 5 – long-pass filter (500 LP, Chroma, USA), 6 – SM-fiber coupler, 7 – SMF-28e fiber, 8 – SSPD, 9 – dichroic mirror (RT 488 rdc).

The excitation of the LG was done using 473 nm picosecond diode laser BDL-473-SMN (Becker&Hickl GmbH, Germany) operating at 50 MHz repetition rate. The average laser power incident on the sample was ∼100 µW. The fluorescence signal from the LG was collected with an objective (LD A-Plan 40x/0.55, Zeiss, Germany), sent to a dichroic mirror (RT 488 rdc)and then coupled to a single-mode fiber (SMF-28e, Corning, US) of the SSPD (Scontel, OPRS-SW-60-wide). A 495 nm long-pass filter (495LP, Chroma, USA) was used to record fluorescence in the desired spectral range. FLIM data was processed using the SPCImage program (Becker&Hickl GmbH, Germany).

The main advantages of SSPDs compared to conventional PMTs or SPADs used in microscopy are high detection efficiency in the broad spectral range both in visible and near-infrared regions, low dark count rates and extremely low jitter [29-31].

The measured system detection efficiency is greater than 60% in a wide wavelength range of 500-950 nm. The operating current of the detector in the experiment was 19 μA, which corresponds to 40 dark counts per second.

## Results and discussion

### Fluorescent properties of LGs before and after their photooxidation. HPLC analysis of LG retinoids

The fluorescent properties of the suspension of native and photooxidized LGs were studied using two different setups: a stationary fluorimeter and a confocal laser scanner setup combined with FLIM and SSPD.

Figure 2A shows the fluorescence spectra of the LG suspension, recorded on a stationary fluorimeter, before (spectrum 1) and after their photooxidation (spectrum 2).

**Fig. 2.**
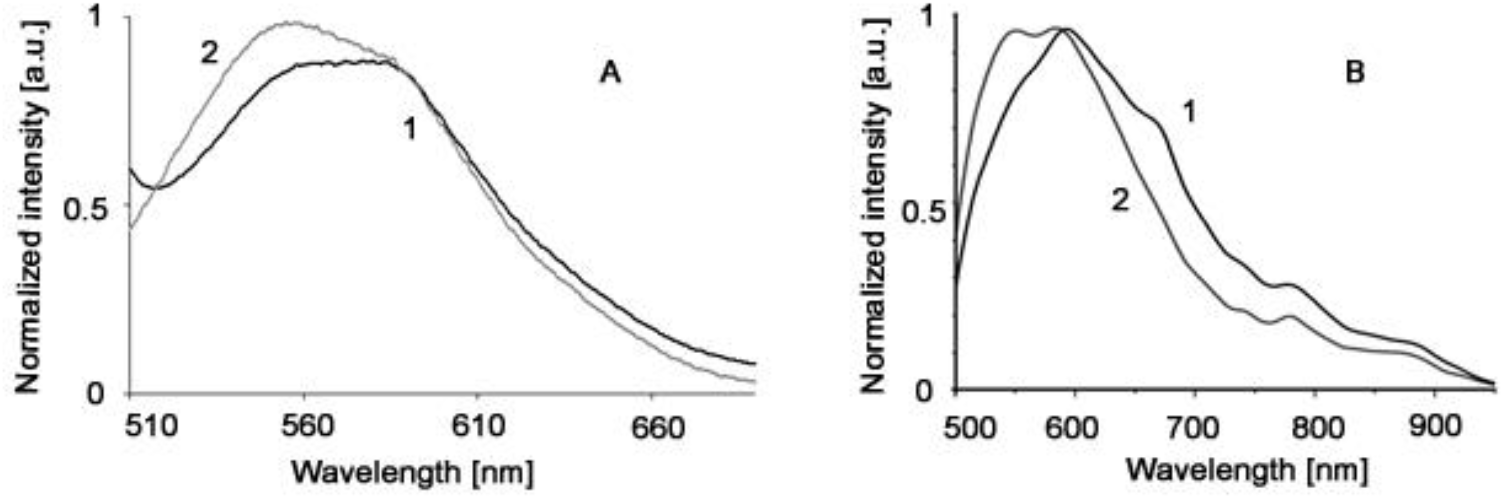
Fluorescence spectra of lipofuscin granule (LGs) suspension: A – normalized spectra obtained on a stationary fluorimeter: 1 – non-irradiated (native) LGs before photooxidation; 2 – LGs photo-oxidized by visible light (400-700 nm). Excitation: 488 nm. B – normalized and fitted spectra obtained using a laser scanner combined with SSPD: 1 – non-irradiated (native) LGs; 2 – LGs photooxidized by UV diode 395 nm, 1 hour. Excitation: 473 nm.

Figure 2B shows the fluorescence spectra of native and photooxidized LG suspensions obtained using a laser scanner combined with SSPD. In these experiments, LG suspension samples in a quartz cuvette were placed in front of the monochromator input in a holder. The radiation was directed at the sample from the side, and the scattered 473 nm laser radiation was cut off by a 500 nm long-pass filter (Chroma) at the monochromator input. The monochromator output was coupled to the optical fiber input of a SSPD detector.

The fluorescence intensity values of the non-oxidized LG sample were measured in the wavelength range of 500-950 nm (Fig. 2B, curve 1). After that, the LG samples was photooxidized with 395 nm radiation at 3 W/cm^2^ for 1 hour. The spectrum for the photooxidized sample is shown in Figure 3B, curve 2

**Fig. 3.**
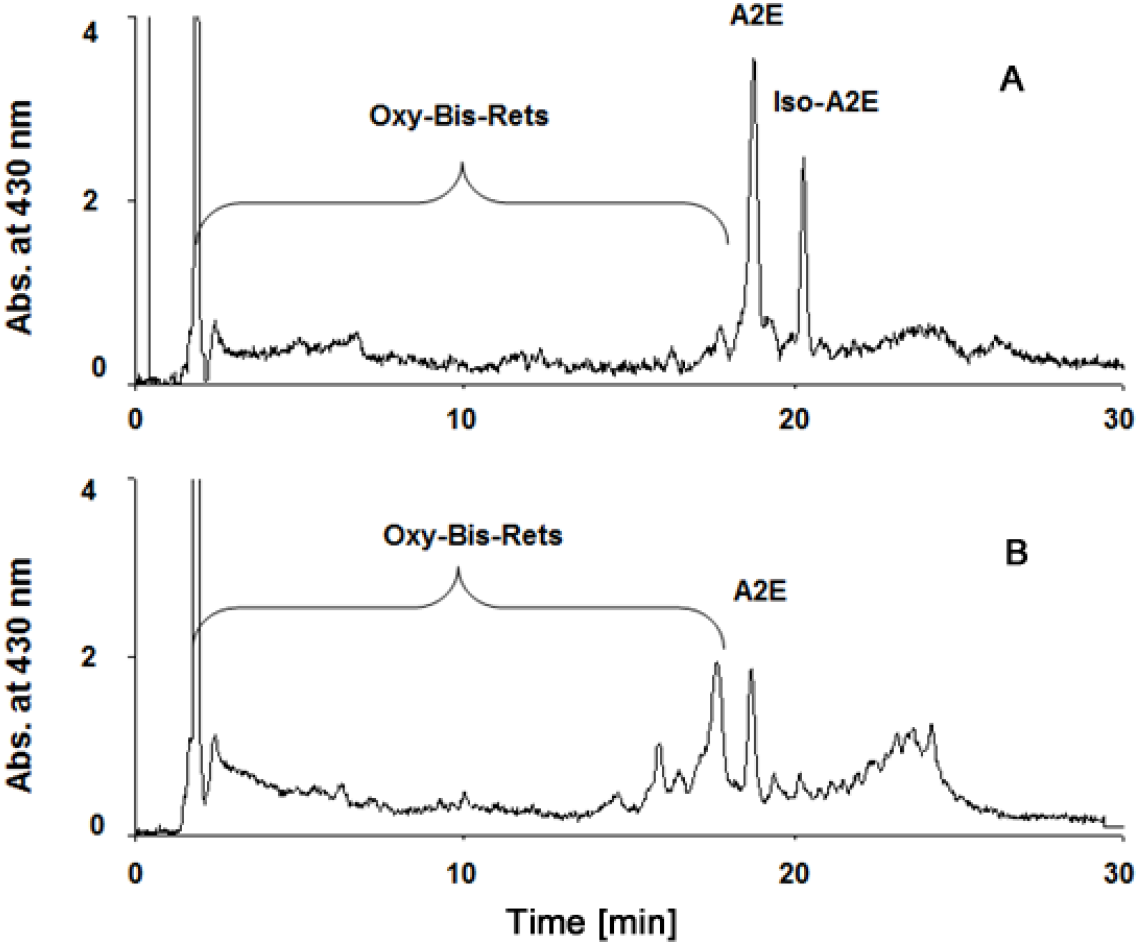
HPLC analysis of chloroform extracts of lipofuscin granules (LGs) containing bisretinoids (Bis-Rets) and products of their photooxidation and photodegradation (Oxy-Bis-Rets): A – native LGs; B – LGs photooxidized by visible light (400-700 nm). Registration at a wavelength of 430 nm.

From the comparison of the fluorescence spectra of native LGs and photooxidized by visible light, obtained using a stationary fluorimeter (Fig. 3A), it is evident that the fluorescence intensity in the short-wave part of the spectrum increases for photo-oxidized LGs. A similar trend is observed in the spectra recorded using a laser scanner combined with SSPD (Fig. 3B), and a redistribution of intensities in the short-wave region of 500-550 nm and the long-wave region of 650-850 nm is also visible for the photooxidized sample.

These data are in good agreement with the results of HPLC analysis. From Figure 4, it is evident that the number of initial peaks at the initial retention times, which determine the presence of Oxy-Bis-Rets in the analyzed system, becomes greater in photo-oxidized LG samples compared to non-oxidized ones. It is the increased content of Oxy-Bis-Rets in photooxidized LGs that leads to a shift in the fluorescence maximum to the short-wave region of the spectrum and an increase in its intensity [11].

**Fig. 4.**
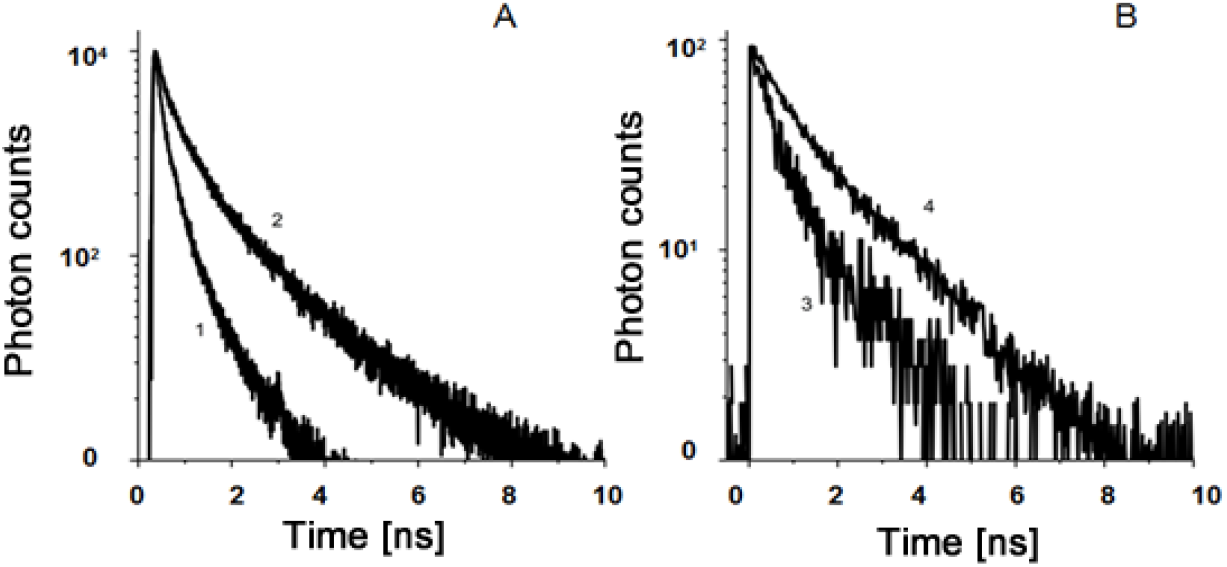
Fluorescence decay curves of lipofuscin granule (LGs) suspension before (1) and after photooxidation (2), approximated by a three-exponential model: A – decay curves recorded on a FluoTime-300 setup (λ_exc_=485 nm, λ_reg_=540 nm); B – decay curves recorded on a FLIM setup (λ_exc_=473 nm, λ_reg_>500 nm).

Thus, comparative fluorescence analysis of native and photooxidized LG samples, performed on a stationary fluorimeter and using a laser scanner combined with SSPD, showed the same tendency of change in the fluorescent properties of LGs during their photooxidation. The difference in the shape of the spectra recorded on different devices may be due to the different conditions of photooxidation of the samples.

### Comparative analysis of the fluorescence decay of native and photooxidized LGs

Previously, the fluorescence decay curves of the LG suspension recorded on a FluoTime-300 fluorimeter (PicoQuant, Germany) were characterized in detail elsewhere [24, 41]. In the present study, similar data were obtained for LG suspension samples before and after their photooxidation on the same setup (Fig. 4A). The fluorescence decay curves of all the samples were recorded at a wavelength of 540 nm with an excitation at a wavelength of 485 nm. Same samples were investigated also by a FLIM setup with SSPD (Fig. 4B).

Using three-exponential model, the c fluorescence lifetimes were calculated for both experiments (Tables 1 and 2).

**Table 1.** Fluorescence lifetimes (τ) of lipofuscin granules (LGs) before and after their photooxidation, obtained using the FluoTime-300 setup (λ_exc_=485 nm, λ_reg_=540 nm) (from the data in Fig. 4A).

| Sample | $\tau_1$, ns | $A_1$ % | $\tau_2$, ns | $A_2$ % | $\tau_3$, ns | $A_3$ % | $\tau_m$ | $A_2+A_3/A_1$ |
| --- | --- | --- | --- | --- | --- | --- | --- | --- |
| native LGs | 0.17 | 66 | 1.1 | 30 | 4.4 | 4 | 0.6 | 0.5 |
| photooxidized LGs | 0.21 | 59 | 1.7 | 34 | 4.9 | 7 | 1.1 | 0.7 |

**Table 2.** Fluorescence lifetimes (τ) of lipofuscin granules (LGs) before and after their photooxidation (λ_exc_=473 nm, λ_reg_>500 nm) using a FLIM system with SSPD (from the data in Fig. 4B).

| Sample | $\tau_1$, ns | $A_1\%$ | $\tau_2$, ns | $A_2\%$ | $\tau_3$, ns | $A_3\%$ | $\tau_m$ | $A_2+A_3/A_1$ |
| --- | --- | --- | --- | --- | --- | --- | --- | --- |
| native LGs | 0.22 | 69.5 | 0.68 | 23.5 | 2.05 | 7 | 0.5 | 0.4 |
| photooxidized LGs | 0.55 | 53.5 | 1.6 | 34.5 | 4.1 | 12 | 1.4 | 0.9 |

Table 1 shows that the contribution of the long-lived component (τ_3_) increases in photo-oxidized LGs (p≤.0.02), while the contribution of the component with the shortest characteristic time (τ_1_) decreases (p .0.04) At the same time, the mean fluorescence lifetime (τ_m_) also increases (p≤0.01), which correlates well with both previous studies and literature data[25, 42, 43].

Previous studieshave shown that Oxy-Bis-Rets have the longest fluorescence lifetimes compared to non-oxidized Bis-Rets[15,24,25]. Fluorescence analysis and HPLC in this work (Figs. 2 and 3) also confirm an increase in the Oxy-Bis-Ret content in the samples during LG photooxidation. Thus, it can be concluded that the increase of τ_3_ and its contribution A_3_ to the fluorescence decay occurs as a result of an increase in the content of Oxy-Bis-Rets in the sample during photooxidation of LGs.

The fluorescence lifetime data (τ_i_) obtained on the FLIM system with SSPD before and after LG photooxidation are presented in Table 2. These values are comparable with the data obtained on the FluoTime-300 setup (Table 1). Despite some discrepancies in the data, which are due to differences in the technical characteristics of the setups, detectors, and data processing programs, as well as the conditions of photooxidation of LG samples, the trend of increasing the contribution of Oxy-Bis-Rets during LG photooxidation is observed in both cases.

Thus, the analysis of the fluorescence decay curves of the native and photooxidized LG suspensions on both setups revealed the same trend: there is an increase in the τ_3_and A_3_ values, that are related to the fluorescence signal from Oxy-Bis-Ret [41]. We would like to point out that τ_2_ and A_2_also increase and thus, they can be the diagnostic markers of the disease.

### Fluorescence lifetime imaging of LGs

Using a FLIM-microscopy system with SSPD, we obtained data on the spatial distribution of characteristic fluorescence lifetimes depending on the spatial coordinates in native and photooxidized LG samples. In contrast to the earlier described point measurements, a detailed visualization of the spatial distribution of fluorescence lifetimes (τ_j_) in LG samples was assessed. The analysis of the data allows observation of the fluorescence lifetime heterogeneity across the LG samples.

Figure 5 shows the data characterizing individual time components (τ_j_) for an array of fluorescence decay curves obtained upon fluorescence excitation of samples containing native and photooxidized LGs. While Figure 5A shows the visualization of the spatial distribution of the fluorescence lifetimes (τ_1_, τ_2_, and τ_3_), figure 5B presents a diagram of the lifetime distribution for each characteristic time (τ_1_, τ_2_, and τ_3_). The data for the native LG sample for each fluorescence lifetime and its contribution (Table 3) correlate well with the point measurements data from the Table 2.

**Table 3.** Fluorescence lifetimes (τ) of lipofuscin granules (LGs) before and after their photooxidation (λ_exc_=473 nm λ_reg_>500 nm) using a FLIM microscopy system and SSPD (from data in Fig. 5B).

| Sample | $\tau_1$, ns | A<sub>1</sub> % | $\tau_2$, ns | A<sub>2</sub>% | $\tau_3$, ns | A<sub>3</sub>% | A<sub>2</sub>+A<sub>3</sub>/A<sub>1</sub> |
| --- | --- | --- | --- | --- | --- | --- | --- |
| native LGs | 0.21 | 65 | 0.69 | 25 | 2.05 | 7 | 0.5 |
| photooxidized LGs | 0.48 and<br>0.95 | 58 | 1.05 and<br>1.7 | 37 | 4.2 and<br>5.2 | 12 | 0.8 |

**Fig. 5.**
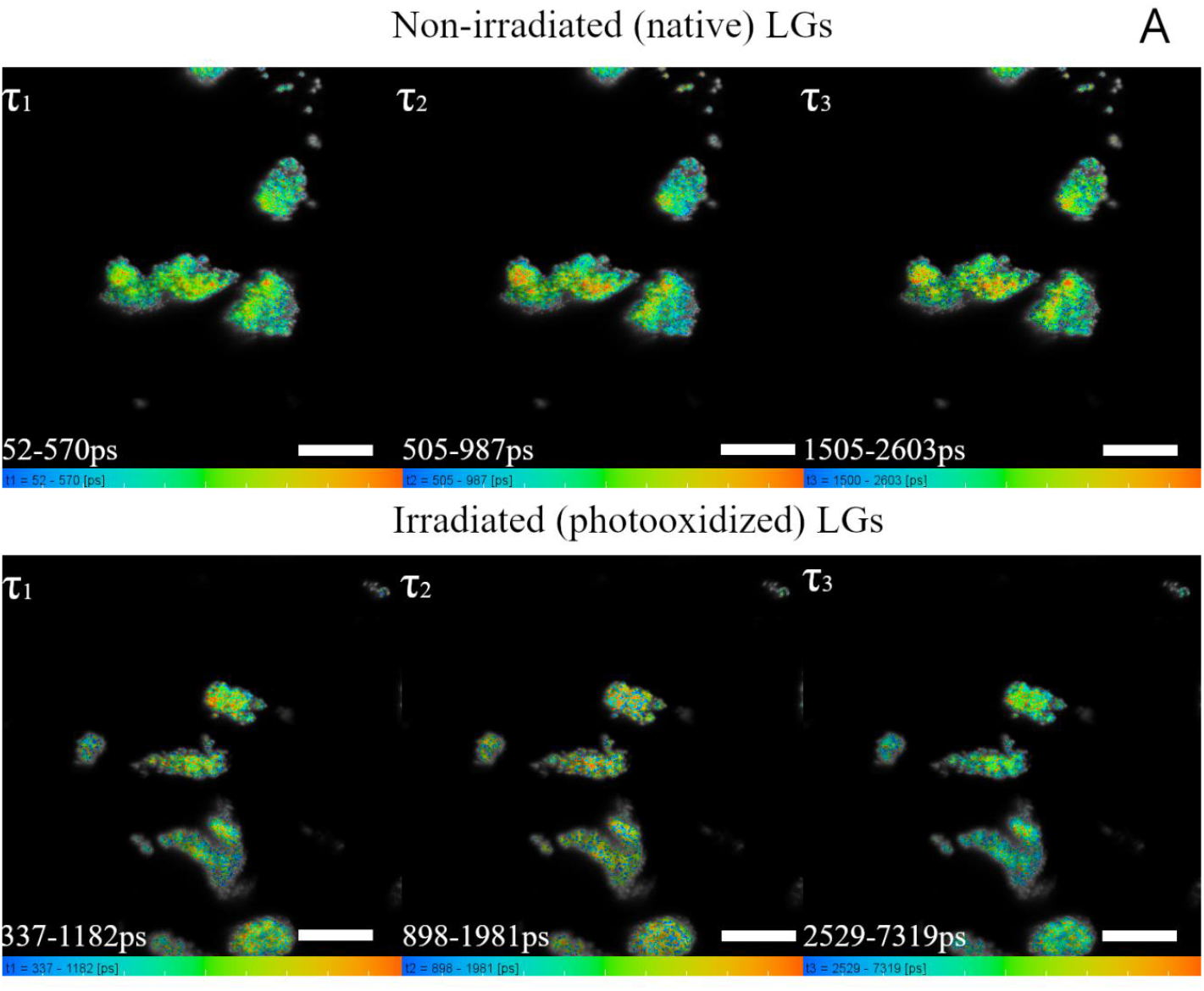
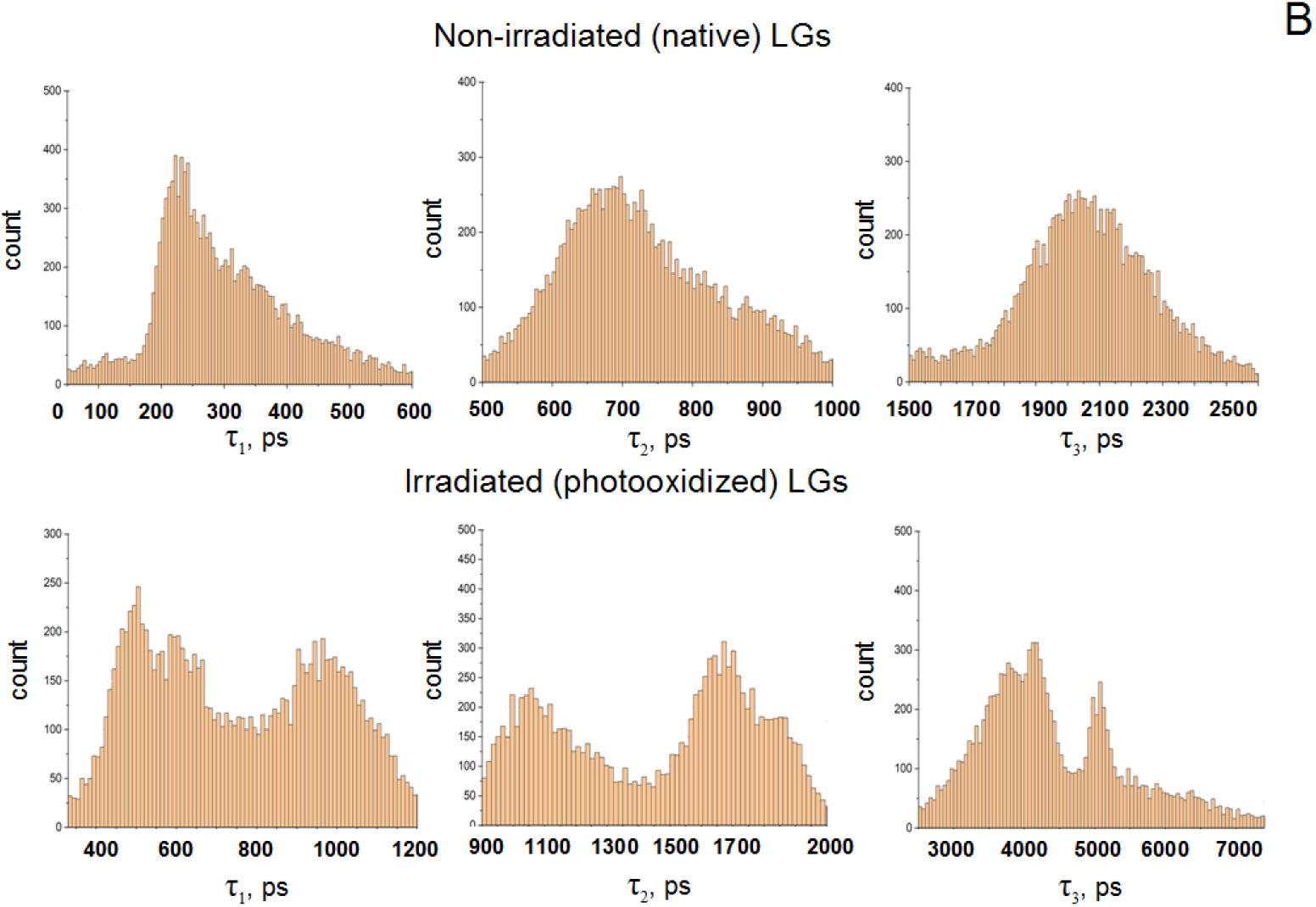
(A) Visualization of the spatial distribution of lifetimes τ_j_ in a lipofuscin granules (LGs) sample before and after photooxidation. Image size: 512×512 pixels (B) Histograms of the distribution of fluorescence lifetimes for each τ_j_ component in samples before and after LG photooxidation.

However, for the samples of photooxidized LGs, such an analysis (Fig. 5B) results in the different from the point measurements data. The histograms show two distinct maxima in the distribution for all characteristic fluorescence lifetimes (τ_1_, τ_2_, and τ_3_), differing from the maximum values for the samples of native LGs. This may indicate that at least two dominant oxidized forms of bisretinoids with quite distinct characteristic fluorescence lifetimes can be produced during the photooxidation of LGs. In any case, for each fluorescence time, an increase in its value is observed in the photooxidized LGs samples compared to the case with native LGs. Table 3 presents maxima of the lifetime values for each τ_i_. However, if we compare the average values of the spatial distribution of fluorescence lifetimes for photooxidized LGs with the data from Table 2, we can note that these values, obtained by different methods on the same setup are very close. Adetailed analysis of the spatial distribution of fluorescence lifetimes for native and photooxidized LGs shows that when searching for numerical criteria of norm and pathology in diagnostics using the FLIM method, using only average values of characteristic fluorescence lifetimes and their contribution to the overall kinetics can lead to incorrect conclusions.

The fact that the fluorescence lifetimes have a broad distribution both for native and photo-oxidized LG samples (Fig. 5B)reflects the significant chemical heterogeneity of LGs caused by the presence of more than 20 different fluorophores[44,45]. However, for the native LG sample, one dominant product can still be identified (Fig. 5B). Photo-oxidized LGs are characterized by a more complex fluorescence lifetime distribution. In the histograms of the photo-oxidized LG sample, more than two dominant products can already be identified. This may be due to the multiplicity of Oxy-Bis-Rets. The set of Oxy-Bis-Rets may depend on the conditions of light exposure (exposure time, light intensity, oxygen level in the system). For example, when irradiating the A2E fluorophore with a molecular weight of 592, oxidized forms of A2E with masses of 608, 624, 640, and 656 can be formed, each of which differs from the previous one in mass by 16 [46]. As the irradiation time increases, the number of oxidized forms of A2E increases relative to the initial non-oxidized form. In addition, we have previously shown that when approximating the LG decay curves using four exponentials, the component with τ_3_, which was represented as a single process in the three-exponential model, is divided into two components. This allows us to identify the presence of both weakly oxidized and strongly oxidized forms of bisretinoids [41]. These data allow us to assume that the histograms of the fluorescence lifetime distribution can change their appearance depending on the sample irradiation conditions. Accordingly, the average values of lifetimes will also vary within a certain range. This must be taken into account when interpreting experimental data and developing diagnostic criteria based on the FLIM method.

To expand the capabilities of the FAF for diagnostics of pathologies the FLIM method is considered a promising approach, but it is necessary to define clear criteria for the normal and pathological states of a tissue.

In addition to the above-described analysis results, visualization of the distribution of amplitudes (A_i_) of fluorescence lifetimes was recorded (Fig. 6). Based on these data, the contributions of each component (τ_1_, τ_2_, and τ_3_) were calculated for the array of decay curves.

**Fig. 6.**
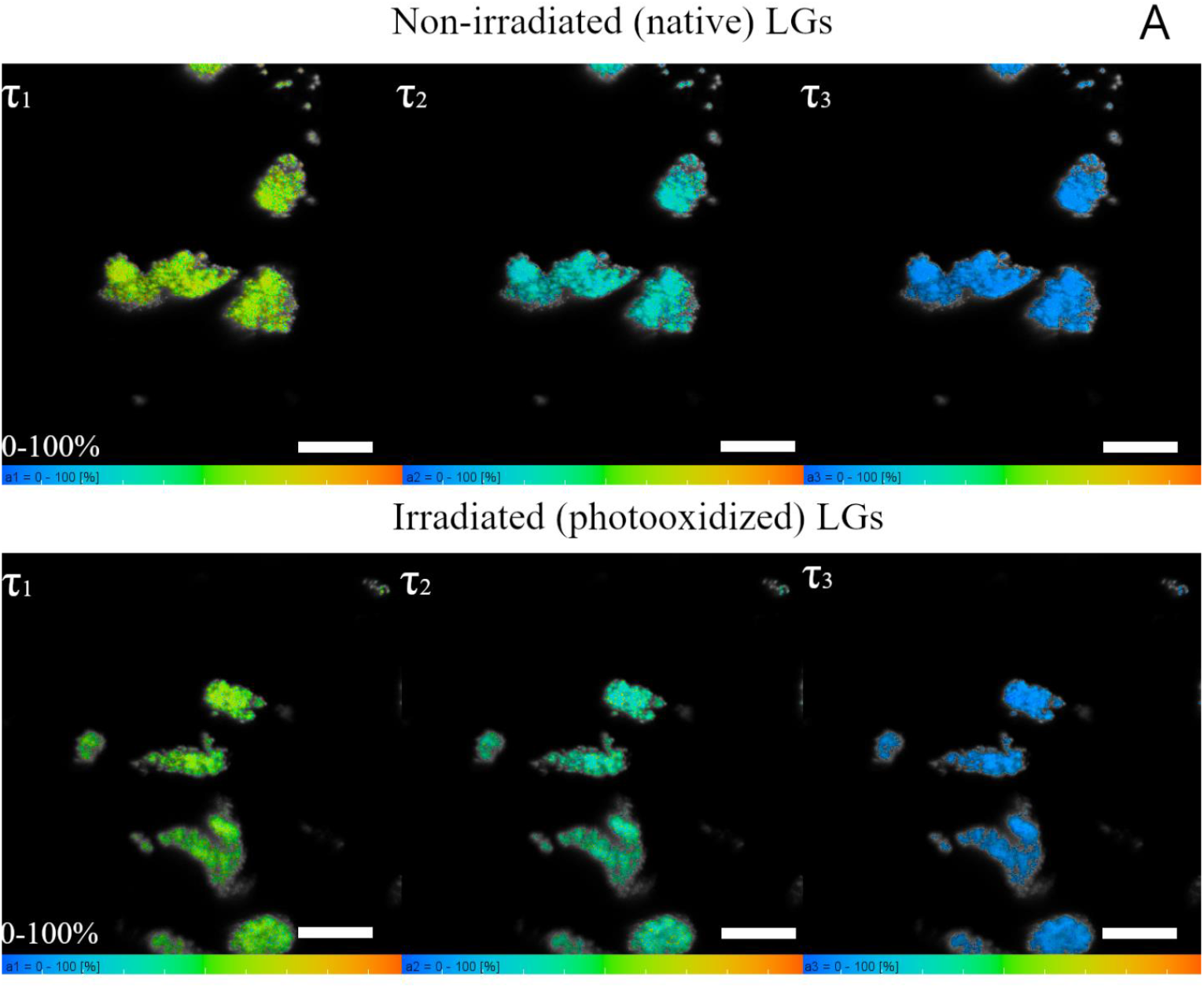
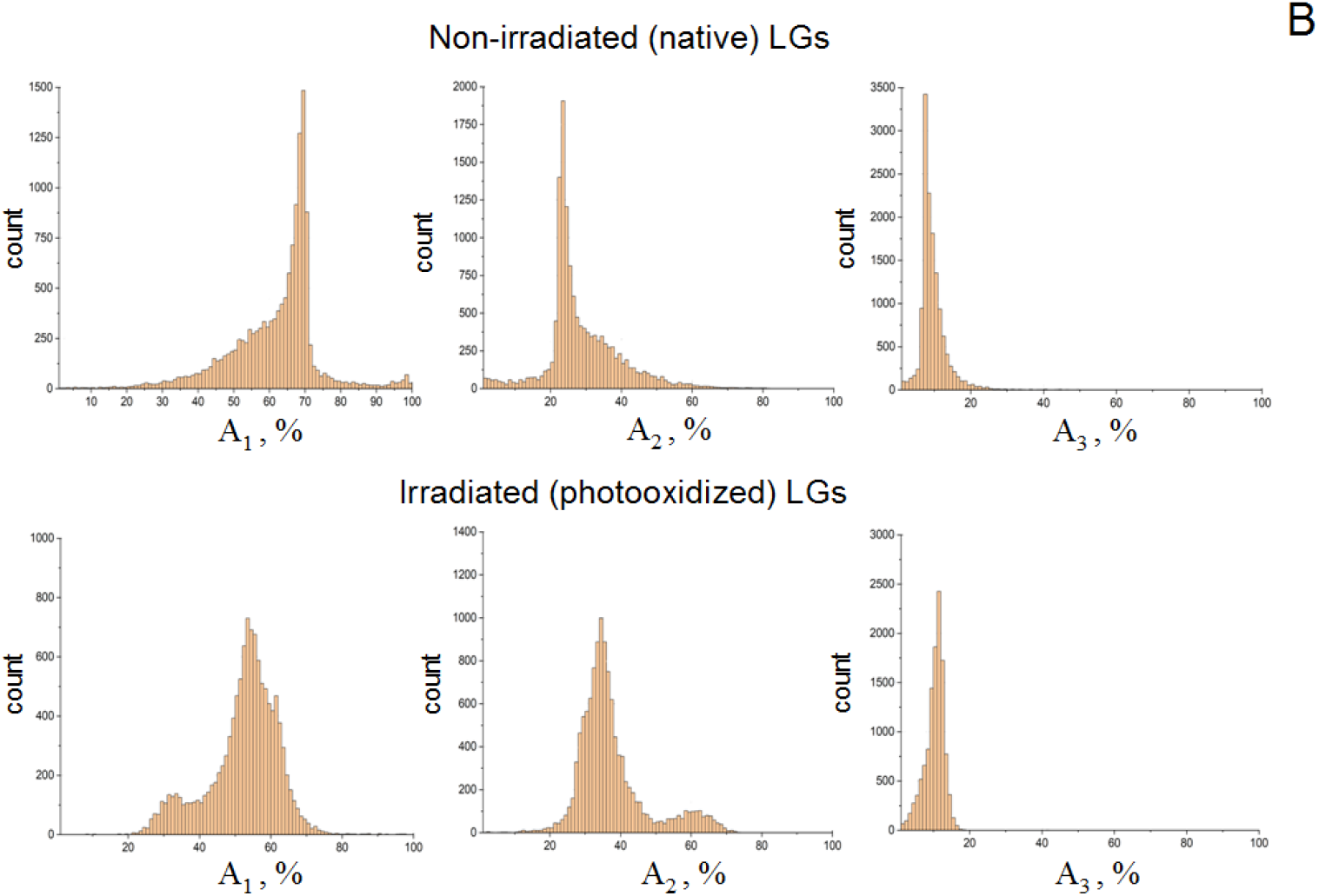
(A) Visualization of the lifetime amplitudes for each of the τ_j_ components (from 0 to 100%) before and after photooxidation of LG samples. Image size: 512×512 pixels. (B) Histograms of the distribution of lifetime amplitudes for each of the τ_j_ components of LGs before and after their photooxidation.

A comparative analysis of the histograms (Fig. 6B) of the lifetime amplitudes for each of the components τ_j_ (Table 3) showed close values of the percentage of dominant products to the averaged data (Table 2) both in native and photooxidiyed LGs. Similar changes in the contribution of the components (τ_1_, τ_2_, and τ_3_) obtained by the FLIM method in two different ways (Figs. 4B and 6A) are observed. This indicates the reliability of the method. During photooxidation, the contribution of the first component (τ_1_) decreases, and the contribution of components with longer times (τ_2_ and τ_3_) increases (Table 3). This analysis does not provide a detailed picture as in the case of histograms of the distribution of fluorescence lifetimes for each τ_j_ component in samples before and after LG photooxidation. Despite slight changes in the shape of the amplitude distribution histograms and the presence of small additional maxima in the case of photooxidized samples, we obtain information similar to that from the averaged analysis. In other words, visualization of fluorescence lifetime amplitudes provides similar information as in the case of the averaged analysis, but does not allow for a deeper differentiation of the stages and degree of photooxidation of LGs.

## Conclusion

In this work, we demonstrate for the first time a laser scanning FLIM system equipped with SSPD for the evaluation of the fluorescence decay parameters of LGs that may have a clinical relevance for the diagnostics of AMD.AMD was simulated in the experiments by the photooxidation of the LGs using visible light. Thus, weevaluated the contributions of Bis-retinoids (Bis-Rets) and Oxidized Bis-retinoids (Oxy-Bis-Rets) to the overall fluorescence decay signal of LGs.

We show that during LG photooxidation, the contribution with the shortest lifetime component decreases, while the contributions of the middle and longest lifetime component increase. Such a change in parameters occurs due to an increase in the Oxy-Bis-Ret content in photooxidized LGs, which in turn leads to an increase in the mean fluorescence decay time. Thus, not only the longest fluorescence lifetimeτ_3_ and its contribution (A_3_) to the overall signal, but also the mean fluorescence lifetime (τ_m_) as well as the other fluorescence lifetimes may be considered valuable parameters in early diagnostics of degenerativeprocesses in the retina and RPE.

## Acknowledgements

We thank Dr. S. Borzenok and Dr. M. Khubetsova from the S. Fyodorov Eye Microsurgery Federal State Institution for their help in contribution of human cadaver eyes.

